# A Neural Network Approach to Identify Left-Right Orientation of Anatomical Brain MRI

**DOI:** 10.1101/2024.02.15.580574

**Authors:** Kei Nishimaki, Hitoshi Iyatomi, Kenichi Oishi

## Abstract

Left-right orientation misidentification in brain MRIs presents significant challenges due to several factors, including metadata loss or ambiguity, which often occurs during the de-identification of medical images for research, conversion between image formats, software operations that strip or overwrite metadata, and the use of older imaging systems that stored orientation differently. This study presents a novel application of deep-learning to enhance the accuracy of left-right orientation identification in anatomical brain MRI scans. A three-dimensional Convolutional Neural Network model was trained using 350 MRIs and evaluated on eight distinct brain MRI databases, totaling 3,384 MRIs, to assess its performance across various conditions, including neurodegenerative diseases. The proposed deep-learning framework demonstrated a 99.6% accuracy in identifying the left-right orientation, thus addressing challenges associated with the loss of orientation metadata. GradCAM was used to visualize areas of the brain where the model focused, demonstrating the importance of the right planum temporale and surrounding areas in judging left-right orientation. The planum temporale is known to exhibit notable left-right asymmetry related to language functions, underscoring the biological validity of the model. More than half of the ten left-right misidentified MRIs involved notable brain feature variations, such as severe temporal lobe atrophy, arachnoidal cysts adjacent to the temporal lobe, or unusual cerebral torque, indicating areas for further investigation. This approach offers a potential solution to the persistent issue of left-right misorientation in brain MRIs and supports the reliability of neuroscientific research by ensuring accurate data interpretation.

## 1. Introduction

The identification of left-right orientation in brain Magnetic Resonance Imaging (MRI) is a crucial step to ensure accurate interpretation and diagnosis [1–5]. Errors in orientation can lead to serious clinical consequences, such as mistaken localization of pathology or even surgical intervention on the wrong side of the brain [2]. For clinical MRI, which is stored in Picture Archiving and Communication Systems (PACS), the MRIs are always displayed in a consistent orientation with digital annotations indicating the left and right sides of the brain in the image viewer. Digital MRIs are usually in Digital Imaging and Communications in Medicine (DICOM) format, which carry metadata that include orientation details. Viewing software can use this information to consistently display images in the correct orientation. As long as we are dealing with DICOM format images, left-right misorientation (or “flipping”) is very unlikely to occur, and when it does, it is usually human error during image manipulation or interpretation. The Neuroimaging Informatics Technology Initiative (NifTI) format is another common format for medical imaging data. The orientation of the data is stored in the header of the NIfTI file, from which many image processing software packages extract the orientation of the MRI data. However, left-right misorientation remains a persistent challenge for several reasons, particularly when dealing with brain MRIs [1].

There can be instances where the orientation information from the header files in DICOM or NIfTI formats is lost or becomes ambiguous. In efforts to de-identify medical images for research or sharing [6, 7], certain metadata in the header might be intentionally stripped, which can sometimes inadvertently include orientation information. Converting medical images from one format to another also can result in a loss of orientation metadata. Some software might strip out or overwrite certain metadata during operations. Older imaging systems might not have stored orientation (or stored it differently) in the header. When such legacy data are integrated with newer systems or software, orientation information might be absent or misinterpreted. Especially in research settings, there might be inconsistency in imaging practices across laboratories or institutions. If proper documentation is not maintained, orientation might become ambiguous. As a result of these causes or combinations of these causes, which is often the case when using multiple image processing software packages sequentially to process the image, the image data can lose their left-right orientation information, which compromises the reliability of neuroscientific research using brain MRI [1].

Once the orientation information is lost from the imaging header, data description is usually the only source of information with which to identify the orientation. To avoid such situations, there have been several attempts to extract orientation information from the image itself, such as attaching fiducial marker to the temple during the scan. Anatomical features of the brain can also help identify the left-right orientation. Typically, the right frontal lobe is slightly more protruded or anterior than the left frontal lobe, and the left occipital lobe is more protruded or posterior than the right occipital lobe, which is called cerebral torque [8–10]. Some of the anatomical structures are known to be asymmetrical, such as the planum temporale [11–13], which is usually larger on the left side compared to the right side. Although these anatomical features can help identify the left-right orientation, there are individual variations and relying on anatomical cues introduces a level of subjectivity, as it depends on the observer’s experience and expertise.

To identify the orientation from the image itself, machine-learning algorithms have been developed to analyze MRI images and detect the orientation, ensuring that the images are displayed correctly. They can also alert clinicians or researchers if a potential left-right flip is detected, although the best reported accuracy remains at 0.96 [14] in two different samples that consisted of 226 and 216 participants, respectively.

The aim of this study was to enhance the accuracy of identifying the left-right orientation of anatomical MRI using deep-learning frameworks. We trained our model on the Alzheimer’s Disease Neuroimaging Initiative (ADNI)-2 dataset and evaluated its accuracy on eight distinct publicly accessible datasets, to assess the model’s performance in cognitively normal adults and individuals with neurodegenerative diseases, including Alzheimer’s disease. The model was named “Laterality Network (LatNet)” and is accessible through the website (URL: https://github.com/OishiLab/LatNet).

## 2. Materials and Methods

### 2.1. Participants

For the training and evaluation of our model, we utilized publicly available brain MRI datasets from several sources: the Alzheimer’s Disease Neuroimaging Initiative 2 and 3 (ADNI2/ADNI3) [15]; the Australian Imaging, Biomarkers and Lifestyle study (AIBL) [16, 17]; the Calgary-Campinas-359 dataset (CC-359) [18]; the LONI Probabilistic Brain Atlas (LPBA40) [19]; the Neurofeedback Skull-stripped repository (NFBS) [20]; and the Open Access Series of Imaging Studies 1 and 4 (OASIS1/OASIS4) [21, 22]. The dataset descriptions utilized in this study are outlined in Table 1. A total of 350 baseline MRIs from ADNI2 were selected for training the LatNet model, alongside an additional 3,384 MRIs obtained from eight distinct brain MRI databases for testing purposes. To mitigate potential bias, only one MRI scan was randomly selected from each participant across these datasets, even though multiple scans might be available for some individuals.

**Table 1.** Dataset used in our study: Alzheimer’s Disease Neuroimaging Initiative 2/3 (ADNI2/ADNI3); Australian Imaging, Biomarkers and Lifestyle (AIBL); Calgary-Campinas-12 (CC-12); LONI Probabilistic Brain Atlas (LPBA40); Neurofeedback Skull-stripped (NFBS) Repository; and Open Access Series of Imaging Studies 1/4 (OASIS1/OASIS4). OASIS4 consists of clinical MRIs with various diseases and conditions. LatNet was trained using only 350 cases in ADNI2. AD, Alzheimer’s disease; MCI, mild cognitive impairment; CN, cognitively normal older people. One MRI per subject was randomly selected to avoid potential bias.

|  |  | Train | Test |  |  |  |  |  |  |  |
| --- | --- | --- | --- | --- | --- | --- | --- | --- | --- | --- |
| Dataset |  | ADNI2 | ADNI2 | ADNI3 | AIBL | CC359 | LPBA40 | NFBS | OASIS1 | OASIS4 |
| Subject |  | 350 | 750 | 929 | 376 | 359 | 40 | 125 | 235 | 570 |
| Spacing (mm) |  | 0.9-1.1 | 0.9-1.3 | 1.0 | 1.0 | 0.9-1.0 | 0.78-0.86 | 1.0 | 1.0 | 0.5-1.1 |
| Slice thickness (mm) |  | 1.2 | 1.2-1.4 | 1.0-1.3 | 1.2 | 1.0-1.3 | 1.5 | 1.0 | 1.2 | 0.9-1.6 |
| Age | mean | 73.7 | 74.3 | 73.7 | 74.0 | 53.4 | 29.2 | 31.0 | 72.3 | 72.5 |
|  | std | 7.4 | 7.5 | 8.0 | 7.1 | 7.8 | 6.3 | 6.6 | 12.0 | 9.2 |
| Sex | F | 160 | 353 | 486 | 198 | 183 | 20 | 77 | 156 | 303 |
|  | M | 190 | 397 | 443 | 178 | 176 | 20 | 48 | 79 | 267 |
| Manufacturer | GE | 97 | 254 | 211 |  | 120 | 40 |  |  |  |
|  | PHILIPS | 65 | 126 | 142 |  | 119 |  |  |  |  |
|  | SIEMENS | 188 | 370 | 576 | 376 | 120 |  | 125 | 235 | 570 |
| Field Strength | 1.5T | 30 | 149 |  | 102 | 179 | 40 |  | 235 | 29 |
|  | 3.0T | 320 | 601 | 929 | 274 | 180 |  | 125 |  | 541 |
| Diagnosis | CN | 122 | 259 | 522 | 268 | 359 | 40 | 125 | 135 | Real-World MRI |
|  | MCI | 196 | 318 | 301 | 64 |  |  |  |  |  |
|  | AD | 32 | 173 | 106 | 44 |  |  |  | 100 |  |

The ADNI [15] was launched in 2003 as a public-private partnership, led by Principal Investigator Michael W. Weiner, MD. The primary goal of ADNI has been to test whether serial magnetic resonance imaging (MRI), positron emission tomography (PET), other biological markers, and clinical and neuropsychological assessment can be combined to measure the progression of mild cognitive impairment (MCI) and early Alzheimer’s disease (AD). For up-to-date information, see www.adni-info.org. The ADNI study has evolved through several phases, with ADNI2 during 2011 - 2016 and ADNI3 during 2016 - 2022 being two of them. In our research, we incorporated Magnetization Prepared Rapid Acquisition Gradient Echo (MPRAGE) images from the ADNI2 and ADNI3 datasets. Following the ADNI team’s guidelines, we selected ADNI2 MPRAGE images of subjects between 55.1 and 94.7 years of age for ADNI2 and subjects 50.5 to 97.4 years of age for ADNI3. These images underwent preprocessing treatments, such as Gradwarp, B1 non-uniformity, and N3 bias field corrections. It is important to note that ADNI3 MPRAGE images did not require this preprocessing as the corrections are automatically applied by the vendors.

The AIBL study, initiated in 2006, focuses on identifying biomarkers and cognitive traits linked to the development of AD. We included original MPRAGE images from AIBL, spanning subjects 55.0 to 96.0 years of age, and these images are available on AIBL’s official website (https://aibl.org.au/).

The CC359 dataset, an open collection of MRIs from healthy adults, utilized scanners from three different vendors (Siemens, Philips, and General Electric (GE)) across magnetic strengths of 1.5 T and 3 T. For our project, we used MPRAGE and 3D spoiled gradient echo sequence (SPGR) images from this dataset, with the SPGR images (subjects 29.0 to 80.0 years of age) sourced from GE (https://www.ccdataset.com/download).

LPBA40 comprises 40 MRI scans of healthy young adults ranging from 19.3-39.5 years of age, using a single 1.5T GE scanner. Our study utilized 3D SPGR images from LPBA40, which were obtained from the website (https://www.loni.usc.edu/research/atlas_downloads).

The NFBS repository provides MRIs of 125 individuals, including 66 diagnosed with various psychiatric disorders, all scanned with a single 3T Siemens scanner. We included MPRAGE images from NFBS, with subjects from 21.0 to 45.0 years of age, in our analysis (http://preprocessed-connectomes-project.org/NFB_skullstripped/).

Last, the OASIS dataset, which offers a wide range of MRI data with which to study normal aging and AD across its four releases (OASIS-1 through OASIS-4), was also part of our study. OASIS4, specifically, is a clinical cohort of subjects who underwent thorough clinical evaluations. We chose MPRAGE images from OASIS, with subjects from 33.0 to 96.0 years of age for OASIS1 and 37.0 to 94.0 years of age for OASIS4, available on the website (https://www.oasis-brains.org/).

### 2.2. Preprocessing

Two distinct datasets were created: images with the correct left-right orientation (Original) and images with the left-right orientation reversed (Flipped). Subsequently, N4 bias field correction [23] was applied to all images to correct intensity non-uniformities. The rationale behind creating the Flipped dataset was to prevent the model from learning the left-right orientation based on the characteristics of the skull-stripping methods, which is discussed in the following section.

### 2.3. Skull-stripping

The goal of this model is to detect images with left-right inversion in T1-weighted whole head scans. Some images in the databases include a fiducial marker to indicate the left-right orientation, such as a vitamin E or fish oil capsule placed on the right temple in the ADNI2 dataset and on the left temple in the OASIS1 dataset. To minimize the risk of the model learning the left-right orientation from the placement of the fiducial marker, skull-stripping was performed prior to model training.

For training images, a deep-learning-based skull-stripping model implemented in OpenMAP-T1 [24] was used. For testing, all images, both Original and Flipped, underwent skull-stripping using two different methods: one based on OpenMAP-T1, and the other on HD-BET, a deep-learning-based method developed with the extensive EORTC-26101 dataset from 37 facilities. Consequently, two types of test images were generated based on the skull-stripping method used: one for OpenMAP-T1 and another for HD-BET. These two distinct test images were created to assess the impact of the skull-stripping method on the accuracy of identifying left-right orientation.

Details of the procedure for creating training and test images are shown in Figure A in the Supplementary Material.

### 2.4. Rigid coregistration to MNI space

After skull-stripping, all images were aligned to the MNI space using a six-parameter rigid transformation in the ANTsPy (https://antspy.readthedocs.io/en/latest/) library. This process involved only rotation and translation for linear transformation, meaning that the size of the brain remained constant, and only the position corrected. The rigid transformations produced images with a voxel size of 2mm x 2mm x 2mm and a matrix size of 80 x 112 x 80. Pixels with intensity values less than 0 or greater than u+3σ (where u is the mean and σ is the standard deviation) were considered outliers and excluded; then, the data was linearly normalized to a range between -1 and 1. The excluded pixels were replaced with the minimum and maximum values of this range.

### 2.5. Model design

The task of detecting incorrect lert-right orientation of the brain is equivalent to classifying images based on whether the left-right orientation is flipped or not. To address this, we developed a neural network model called LatNet specifically for identifying left-right orientation flips. The LatNet is a 3D Convolutional Neural Network (3D-CNN) comprising four blocks that include convolution layers, batch normalization, ReLU activation functions, and max pooling layers, followed by two fully connected layers. Figure 1 illustrates the architecture of the LatNet.

**Figure 1.**
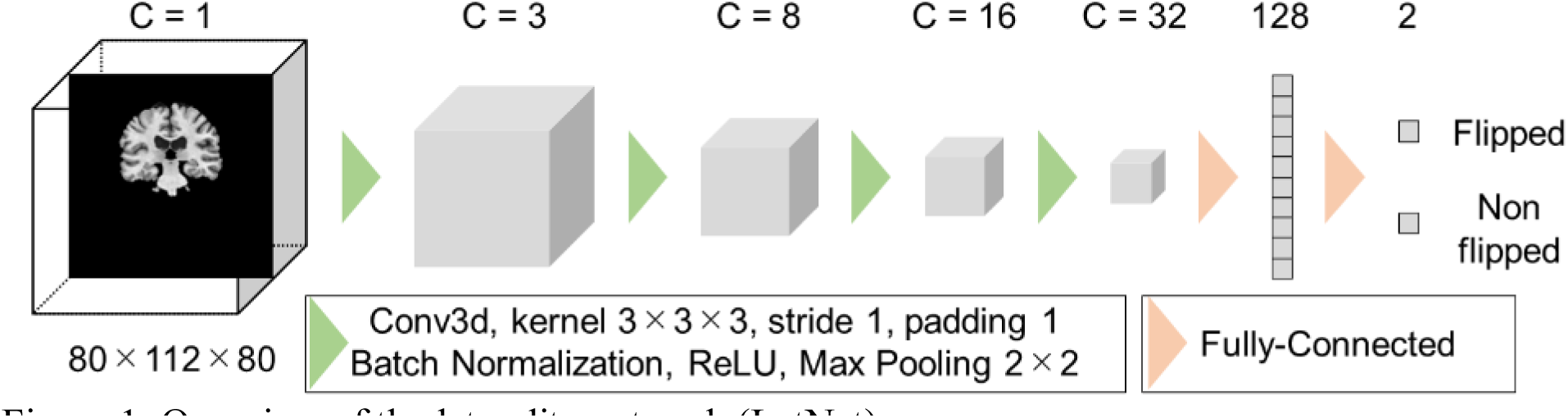
Overview of the laterality network (LatNet).

LatNet is designed to take skull-stripped brain MRI images as input and determine whether the brain has been flipped. During the training phase of the model, a randomized method was employed in which each brain MRI image had a 50% chance of being horizontally flipped. Images that were not flipped were labeled as ’0,’ and those that were flipped were labeled as ’1.’ As this is a binary classification problem, cross-entropy was chosen as the loss function.

To enhance the reliability of the prediction, LatNet used the average prediction probability of models constructed with five different seed values to generate its final results. This methodology is designed to mitigate the variability that can arise from the initialization weight, ensuring a more stable and dependable performance across multiple runs [25].

Furthermore, to ensure the model identifies brain flips based on anatomical structure rather than extraneous factors, such as the position or angle of the brain, we introduced random rotations between -10 to +10 degrees and random translations between -10 to +10 pixels across all axes with a 100% probability during training. This ensured that the brain’s position and angle varied completely randomly in the training of the LatNet.

The LatNet was trained on a single RTX 3090 GPU with 24GB of memory for approximately 24 hours. Automatic Mixed Precision (AMP) technology was used to accelerate the training process. The training was conducted over 10,000 epochs, with a learning rate that gradually decreased from 0.01 to 0.0001, following a Cosine Annealing Learning Rate Scheduler. The batch size was set to 64.

### 2.6. Evaluation of identification performance

An accuracy score was utilized to assess the performance of the LatNet in identifying the correct laterality. Note that for each case, both Original and Flipped images were provided, resulting in a label ratio of one-to-one. Therefore, it is sufficient to use only accuracy as the evaluation metric, which reflects the ratio of correctly identified cases of left-right inversion to the total cases evaluated.

To clarify which areas of the brain LatNet prioritizes to distinguish between left and right orientations, Gradient-weighted Class Activation Mapping (Grad-CAM) [26] was used. Grad-CAM is a method that elucidates the reasoning behind the decisions of CNNs in image classification tasks, and generates a heatmap that highlights important regions of an input image. Grad-CAM examines the gradients flowing into the final convolutional layer of the CNN in image classification tasks by generating a heatmap that accentuates the significant regions of the input image. It does this by analyzing the gradients that flow into the final convolutional layer of the CNN, determining the importance of each neuron for the prediction of a particular class. These importance levels are then combined in a weighted manner and mapped back onto the input image to create a heatmap. This heatmap visually demonstrates the parts of the image that were deemed crucial by the CNN, providing insight into the areas the network finds most relevant for classifying a specific orientation.

### 2.7. Ethics statement

This study did not involve the collection of new data; instead, it utilized exclusively previously published image datasets listed in 2.2. Participants. These datasets were anonymized to safeguard the privacy and confidentiality of the individuals featured. The acquisition of these images adheres to ethical guidelines, with approvals obtained from the appropriate Institutional Review Boards (IRBs).

## 3. Results

Table 2 presents the accuracy of the LatNet in detecting laterality errors across eight datasets, as well as the count of misclassified cases. The accuracy scores are derived from the average probabilities of models created using five distinct seed values. The average accuracy and standard deviation for models created with each of the five seed values are presented in Supplementary Material, Table A.

**Table 2.** Accuracy and number of misclassified cases for models created using five different seed values. The accuracy is calculated based on the average predicted probabilities from these models. # failed refers to the number of cases in which the <u>prediction was incorrect, based on the average predicted probability.</u>

| Dataset | # data | OpenMAP-T1 |  |  |  | HD-BET |  |  |  |
| --- | --- | --- | --- | --- | --- | --- | --- | --- | --- |
|  |  | Original |  | Flipped |  | Original |  | Flipped |  |
|  |  | # failed | Accuracy (%) | # failed | Accuracy (%) | # failed | Accuracy (%) | # failed | Accuracy (%) |
| ADNI2 | 750 | 1 | 99.87 | 0 | 100.00 | 0 | 100.00 | 0 | 100.00 |
| ADNI3 | 929 | 0 | 100.00 | 0 | 100.00 | 0 | 100.00 | 0 | 100.00 |
| AIBL | 376 | 0 | 100.00 | 1 | 99.73 | 0 | 100.00 | 0 | 100.00 |
| CC359 | 359 | 5 | 98.61 | 3 | 99.16 | 2 | 99.44 | 2 | 99.44 |
| LPBA40 | 40 | 0 | 100.00 | 0 | 100.00 | 0 | 100.00 | 1 | 97.50 |
| NFBS | 125 | 0 | 100.00 | 0 | 100.00 | 0 | 100.00 | 0 | 100.00 |
| OASIS1 | 235 | 3 | 98.72 | 2 | 99.15 | 6 | 97.45 | 2 | 99.15 |
| OASIS4 | 570 | 1 | 99.82 | 1 | 99.82 | 1 | 99.82 | 1 | 99.82 |

LatNet demonstrated a 100% accuracy rate in several datasets, including ADNI3, AIBL, LPBA40, and NFBS. The minimum accuracy achieved was 97.45%, which is still considered outstanding. It is important to note that most misclassifications occurred within the CC359, OASIS1, and OASIS4 datasets.

Figure 2 illustrates areas of strong response in the average Grad-CAM across each dataset, highlighting the regions that received the most focus from the model. The average Grad-CAM images for each dataset, corresponding to each seed, are provided in Figures B-F in the Supplementary Material. Notably, all datasets consistently observed a significant response in the right planum temporale and adjacent areas, such as the superior and middle temporal gyri.

**Figure 2.**
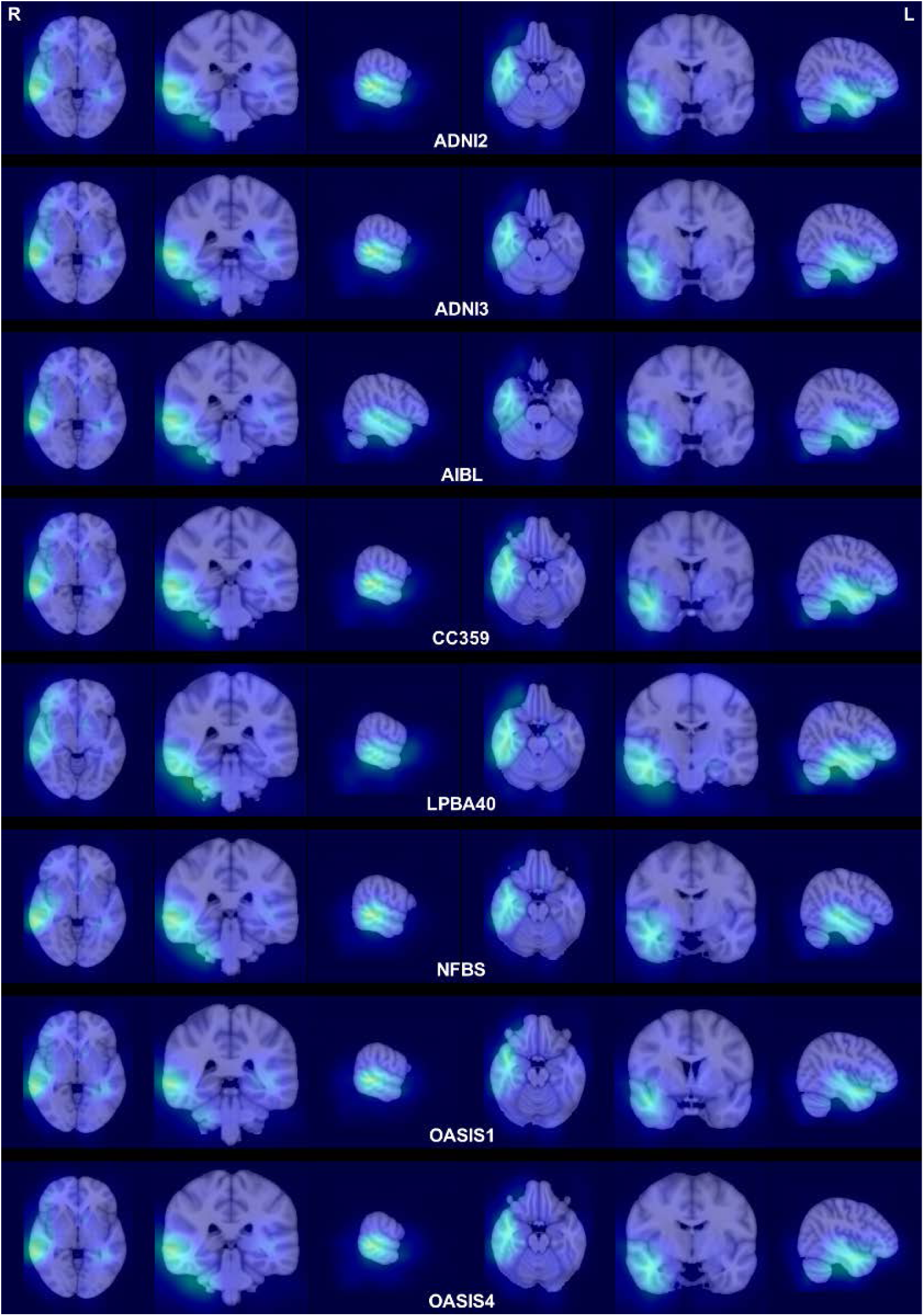
Average Grad-CAM visualizations were generated for images correctly identified with their left-right orientation. Specifically, for each image, an average visualization was created from five Grad-CAM outputs, each produced by a model initialized with a different seed. Subsequently, all images were aligned to the MNI-152 space [27] using the ANTsPy library. Grad-CAM visualizations were then averaged for each dataset within the MNI space.

Figure 3 presents ten instances where the LatNet incorrectly classified the images (3rd column from the left of Table 2), with models based on five different seeds. The left column features Grad-CAM visualizations for correct predictions, whereas the right column includes Grad-CAM visualizations for incorrect predictions. In the correct predictions column, Grad-CAM primarily emphasized the right planum temporale and adjacent regions, similar to the observations in Figure 2. Conversely, the incorrect predictions column predominantly highlighted the left planum temporale and its adjacent areas.

**Figure 3.**
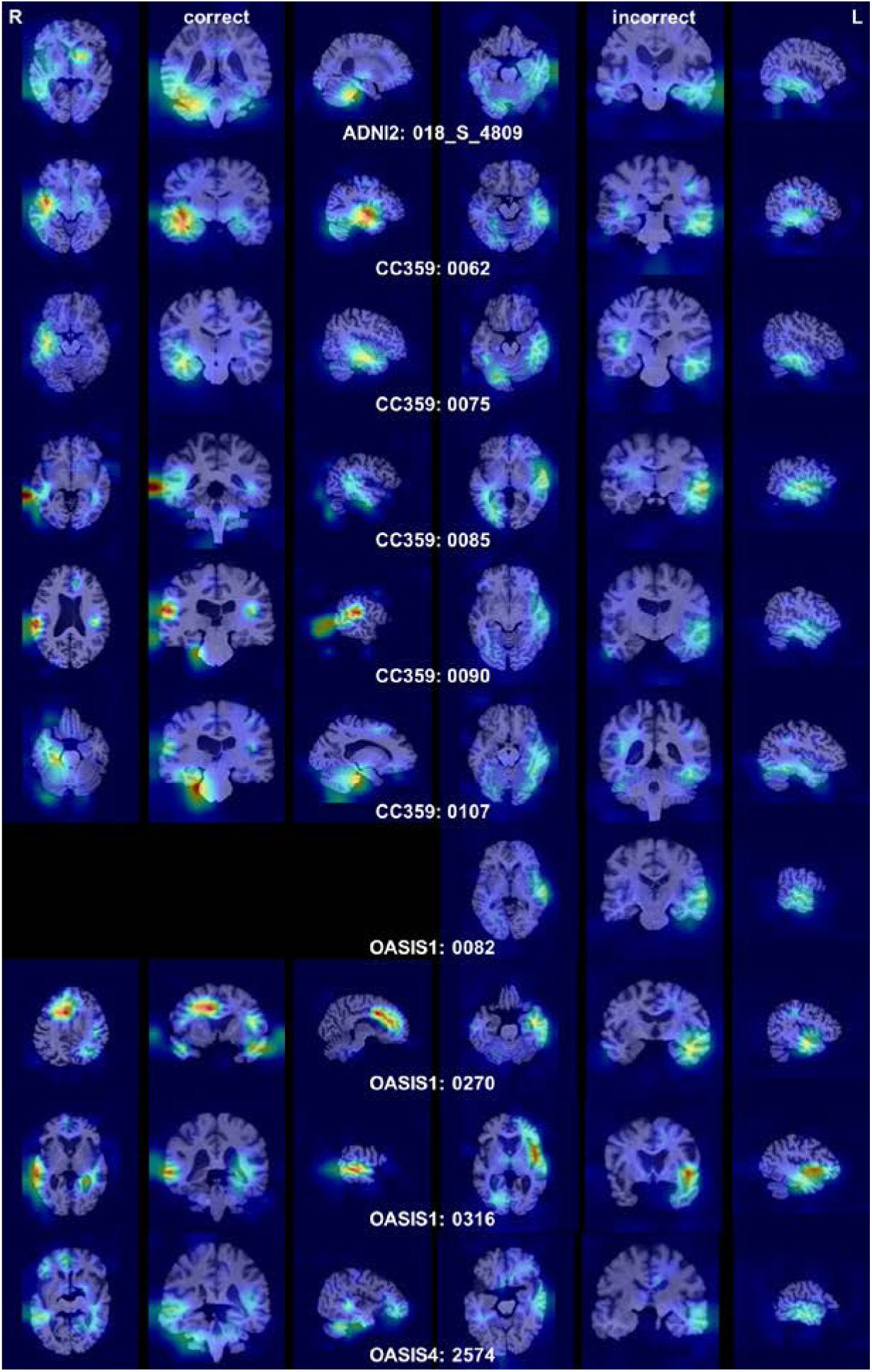
The cross-sections of average Grad-CAM visualizations derived from 10 images with incorrect predictions. This corresponds to the # failed column of the OpenMAP-T1 Original column in Table 2. The left side of Figure 3 displays the average Grad-CAM images from models that accurately identified the left and right orientation amoung the five models, whereas the right side shows the average Grad-CAM images from models that made incorrect predictions amoung the five models. The absence of images in the left column for OASIS1-0082 signifies that all five models failed to predict correctly, as indicated by the blank space. Notably, the anatomical features of CC0090 and CC0107 were remarkably similar, suggesting that these images might have come from the same participant, despite the differing subject IDs.

A closer examination of these ten cases revealed pathological changes or anatomical variations in more than half of them, which are presumed to have led to the incorrect predictions. Specifically, severe temporal lobe atrophy was noted in case 018_S_4809 from the ADNI2 dataset and in case 0316 from OASIS1. An opposite cerebral torque was identified in case 0075 from the CC359 dataset, while ventricular asymmetry was observed in cases 0090 and 0107, and an arachnoid cyst was observed adjacent to the right temporal pole in case 2574 from OASIS4. However, the remaining images did not display any distinctive features that could explain the reasons for the incorrect predictions.

## 4. Discussion

This study presents the creation and application of a deep-learning model capable of distinguishing between the left and right sides of brain MRI images. The model was applied to 3,384 original images and to the same number of horizontally flipped images across eight types of test data, 7,468 in total. Due to the use of fiducial markers in some databases to prevent left-right confusion, the model was trained on images processed with skull-stripping. To prevent the neural network from learning the left-right features of the brain from brain masks obtained through skull-stripping, the model was tested on images that were skull-stripped using a different method (HD-BET) than the one used in the training data (OpenMAP-T1). The model achieved high accuracy rates on all test data, correctly identifying the left and right sides in 99.8%, regardless of the skull-stripping methods.

An existing tool for detecting image left-right flip has the capability to identify discrepancies in the left-right orientation within pairs of images of the same individual, such as between T1-weighted and echo planar images . This tool was tested on 90 pairs of images with mismatched orientations and 88 pairs with matched orientations, and has been reported to accurately determine orientation concordance and discordance with 100% accuracy [1]. However, this tool is capable of detecting only inconsistencies between paired images in terms of left and right orientation. As an attempt to distinguish between the left and right hemispheres of the brain solely based on anatomical information, there is a method by [14] that utilizes machine-learning. Those authors created a model with which to distinguish between the right hemisphere and the left hemisphere with approximately 97% accuracy, based on the morphological information from each voxel, using Least Absolute Shrinkage and Selection Operator on images of the right hemisphere and left-right flipped images of the left hemisphere. However, to our knowledge, there are no existing open-source tools capable of determining the left and right sides of T1-weighted MRI images without pre-processing, making our developed model the first of its kind. Its accuracy significantly surpasses that of the existing machine-learning model and is likely to become a benchmark for the development of future tools for left-right determination.

Using GradCAM, we explored the brain regions the model referenced to determine left-right orientation. The model predominantly focused on the right planum temporale, known for its distinct left-right asymmetry, typically larger on the left due to its association with language functions [11–13]. Interestingly, the planum temporale is also highlighted as a particularly important structure for the differentiation between the left and right sides in the machine-learning model developed by [14]. A previous report that utilized a data-driven approach based on voxel-based analysis identified regions within the brain structure that exhibited significant left-right volumetric differences. The planum temporale, as well as the superior occipital gyrus (larger on the left than the right), the frontal lobe (larger on the right than the left), the anterior insula (larger on the right than the left), and the head of the caudate nucleus (larger on the right than the left), has been recognized as one of the areas with the most pronounced left-right volume asymmetry [28]. Moreover, it is known that the cerebrum exhibits a morphological characteristic known as cerebral torque, which is the tendency of the right hemisphere to rotate slightly forward relative to the left [8–10]. This may result in a larger and wider right frontal lobe, and a wider left occipital lobe that protrudes rightward. This torque is thought to cause the left Sylvian fissure to be more extended than the right, consequently making the left planum temporale larger than its right counterpart. These neuroscientific findings support the biological validity of our model. Despite the fact that the prevalence of planum temporale asymmetry is found to be around 65% [29], our model was able to identify the left and right sides with near 100% accuracy. This suggests that deep-learning might be capable of extracting information beyond traditional quantitative measures, such as volume and surface area.

There were ten images in which left-right orientation was incorrectly identified using the average prediction probability from five models. Due to the small number of these erroneous cases, it was difficult to statistically identify the reasons for misidentification. Analysis using GradCAM revealed that all of the misidentified cases showed a response pattern focused on areas around the left planum temporale, which is opposite to the correct cases. Upon detailed examination of these images, half exhibited clear visual characteristics, including notable atrophy in the temporal lobe, including in the planum temporale, pronounced asymmetry in the volume of the ventricles, and large subarachnoid cysts that displaced the anterior part of the right temporal lobe. Furthermore, there were cases with non-pathological variations, such as an unusual cerebral torque where the left frontal lobe protruded more anteriorly than the right, and the right occipital lobe extended further posteriorly than the left. However, some misidentifications occurred without clear pathological changes or variations in cerebral torque, indicating a need for further investigation.

The model created has a high accuracy rate for determining left-right orientation, suggesting its potential use for screening large volumes of images to check for any left-right reversals. However, since the accuracy is not perfect, additional measures, such as verifying the headers of the original DICOM data, are necessary for images suspected of left-right reversal.

## 5. Limitations

This model has several limitations. Notably, some misidentified images exhibited significant atrophy in the temporal lobe or conspicuous asymmetry in the volume of the ventricles, indicating the need for caution when dealing with images with severe brain atrophy or asymmetry. The accuracy rate of the model on the ADNI data was high, suggesting robustness against morphological changes induced by Alzheimer’s disease. However, in clinical practice, more severe cases than those found in the ADNI cohort might be subject to analysis, necessitating verification of the extent to which the model can tolerate changes. There is evidence to suggest that the asymmetry in the volume of the planum temporale may be less pronounced in schizophrenia. Moreover, studies have observed a reduced volume of gray matter in the left planum temporale in schizophrenic and dyslexic [29–32] patients, with a reversal of the typical asymmetry where the left is larger than the right. Therefore, it is necessary to validate whether this model can accurately determine left-right orientation in brain MRIs of schizophrenic patients. If the model does misclassify the left-right orientation in schizophrenic patients or dyslexic patients, there may be a new potential diagnostic use for this model.

## 6. Conclusion

This study developed a tool capable of determining the left-right orientation of T1-weighted MRI images with over 99.6% accuracy using only anatomical information. The model primarily focuses on the asymmetry of the temporal plane for lateralization determination. Future investigations are required to assess the model’s accuracy in the presence of brain deformities or atrophy due to various diseases.

## Data and Code Availability

The source code of LatNet is available at: https://github.com/OishiLab/LatNet. Each dataset used in this paper is available at www.adni-info.org for ADNI2/3, https://aibl.org.au/ for AIBL, https://www.ccdataset.com/download for CC359, https://www.loni.usc.edu/research/atlas_downloads for LPBA40, http://preprocessed-connectomes-project.org/NFB_skullstripped/forNFBS, and https://www.oasis-brains.org/ for OASIS1/4.

## Author Contributions

Kei Nishimaki: Data curation, Formal analysis, Investigation, Methodology,Software, Validation, Visualization, Writing – original draft. Hitoshi Iyatomi: Resources, Writing – review & editing. Kenichi Oishi: Conceptualization, Project administration, Supervision, Writing – review & editing.

## Acknowledgements

The MRI data collection and sharing for this project was funded by the Alzheimer’s Disease Neuroimaging Initiative (ADNI) (National Institutes of Health Grant U01 AG024904) and DOD ADNI (Department of Defense award number W81XWH-12-2-0012). ADNI is funded by the National Institute on Aging, the National Institute of Biomedical Imaging and Bioengineering, and through generous contributions from the following: AbbVie; Alzheimer’s Association; Alzheimer’s Drug Discovery Foundation; Araclon Biotech; BioClinica, Inc.; Biogen; Bristol-Myers Squibb Company; CereSpir, Inc.; Cogstate; Eisai Inc.; Elan Pharmaceuticals, Inc.; Eli Lilly and Company; EuroImmun; F. Hoffmann-La Roche Ltd. and its affiliated company Genentech, Inc.; Fujirebio; GE Healthcare; IXICO Ltd.; Janssen Alzheimer Immunotherapy Research & Development, LLC.; Johnson \& Johnson Pharmaceutical Research & Development LLC.; Lumosity; Lundbeck; Merck & Co., Inc.; Meso Scale Diagnostics, LLC.; NeuroRx Research; Neurotrack Technologies; Novartis Pharmaceuticals Corporation; Pfizer Inc.; Piramal Imaging; Servier; Takeda Pharmaceutical Company; and Transition Therapeutics. The Canadian Institutes of Health Research is providing funds to support ADNI clinical sites in Canada. Private sector contributions are facilitated by the Foundation for the National Institutes of Health (www.fnih.org). The grantee organization is the Northern California Institute for Research and Education, and the study is coordinated by the Alzheimer’s Therapeutic Research Institute at the University of Southern California. ADNI data are disseminated by the Laboratory for Neuro Imaging at the University of Southern California.

Data were provided by OASIS1/4: Cross-Sectional: Principal Investigators: D. Marcus, R. Buckner, J. Csernansky, J. Morris; P50 AG05681, P01 AG03991, P01 AG026276, R01 AG021910, P20 MH071616, U24 RR021382 and Clinical Cohort: Principal Investigators: T. Benzinger, L. Koenig, P. LaMontagne.

