## Supplementary Table 1 for "A Neural Network Approach to Identify Left-Right Orientation of Anatomical Brain MRI"

### SUPPLEMENTARY MATERIALS

Table A. The average and standard deviation of accuracy and misclassified cases for each model created using five different seed values.

| | | OpenMAP-T1 | | | | HD-BET | | | |
|---|---|---|---|---|---|---|---|---|---|
| | | Original | | Flipped | | Original | | Flipped | |
| Dataset | # data | # failed | Accuracy (%) | # failed | Accuracy (%) | # failed | Accuracy (%) | # failed | Accuracy (%) |
| ADNI2 | 750 | 3.0 (±6.9) | 99.60 (±0.2) | 2.4 (±6.2) | 99.68 (±0.83) | 3.4 (±5.4) | 99.55 (±0.73) | 2.0 (±6.7) | 99.73 (±0.89) |
| ADNI3 | 929 | 2.4 (±6.1) | 99.74 (±0.1) | 1.4 (±3.0) | 99.85 (±0.32) | 4.4 (±6.9) | 99.53 (±0.75) | 1.2 (±2.2) | 99.87 (±0.24) |
| AIBL | 376 | 0.6 (±1.0) | 99.84 (±0.1) | 0.6 (±1.0) | 99.84 (±0.25) | 0.2 (±0.9) | 99.95 (±0.23) | 0.4 (±1.8) | 99.89 (±0.47) |
| CC359 | 359 | 5.2 (±9.0) | 98.55 (±0.5) | 3.4 (±5.1) | 99.05 (±1.41) | 6.0 (±5.8) | 98.33 (±1.60) | 6.2 (±11.9) | 98.27 (±3.33) |
| LPBA40 | 40 | 0.2 (±0.5) | 99.50 (±0.5) | 2.4 (±4.1) | 94.00 (±10.18) | 1.4 (±2.8) | 96.50 (±6.91) | 3.8 (±7.8) | 90.50 (±19.57) |
| NFBS | 125 | 0.6 (±1.7) | 99.52 (±1.4) | 0.0 (±0.0) | 100.00 (±0.00) | 0.4 (±1.0) | 99.68 (±0.83) | 0.0 (±0.0) | 100.00 (±0.00) |
| OASIS1 | 235 | 7.8 (±16.5) | 96.68 (±7.0) | 4.4 (±8.4) | 98.13 (±4.40) | 12.8 (±17.4) | 94.55 (±7.41) | 5.2 (±7.7) | 97.79 (±3.26) |
| OASIS4 | 570 | 1.6 (±2.7) | 99.72 (±0.5) | 3.2 (3.1) | 99.44 (±0.54) | 3.4 (±3.0) | 99.40 (±0.53) | 3.8 (±7.1) | 99.33 (1.24) |
