## Supplementary Figure A for "A Neural Network Approach to Identify Left-Right Orientation of Anatomical Brain MRI"

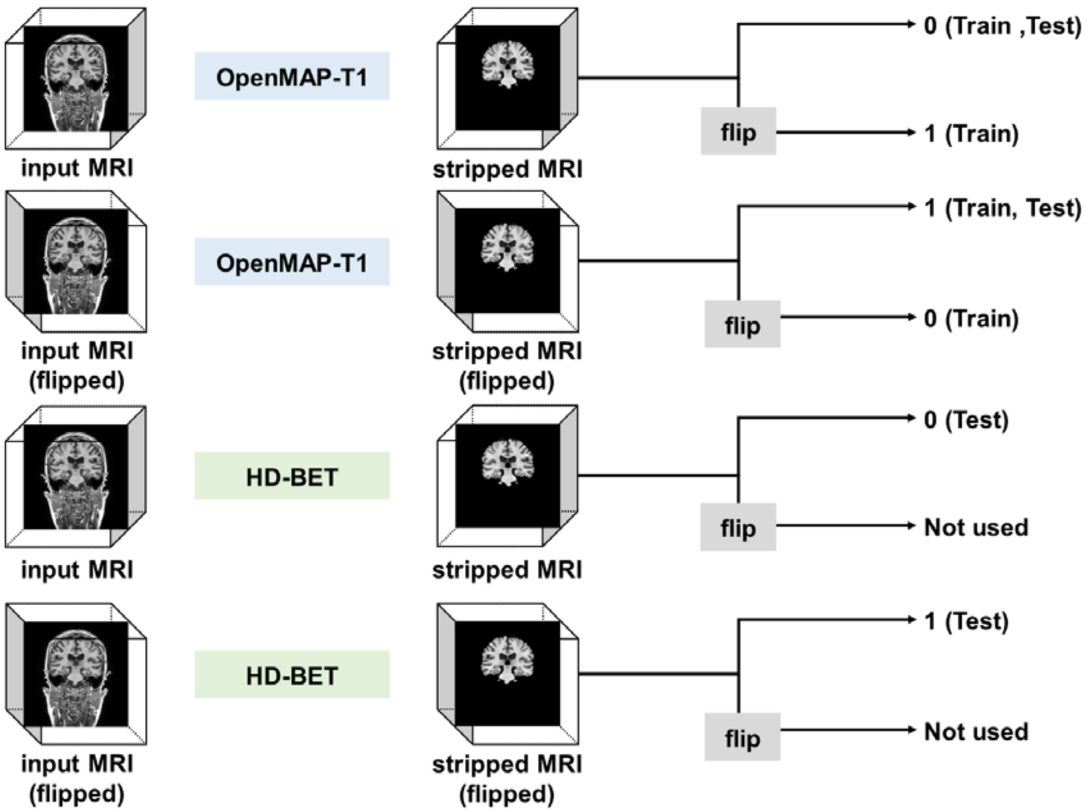


Supplementary Figure A. Overview of the skull-stripping. OpenMAP-T1 applied skull-stripping to the original MRI non-flipped and flipped images, but HD-BET was applied only to the non-flipped images. The 0 indicates a non-flipped image, and the 1 indicates a flipped image.
