## Supplementary figures and images for "A Neural Network Approach to Identify Left-Right Orientation of Anatomical Brain MRI"

### Supplementary Figure B

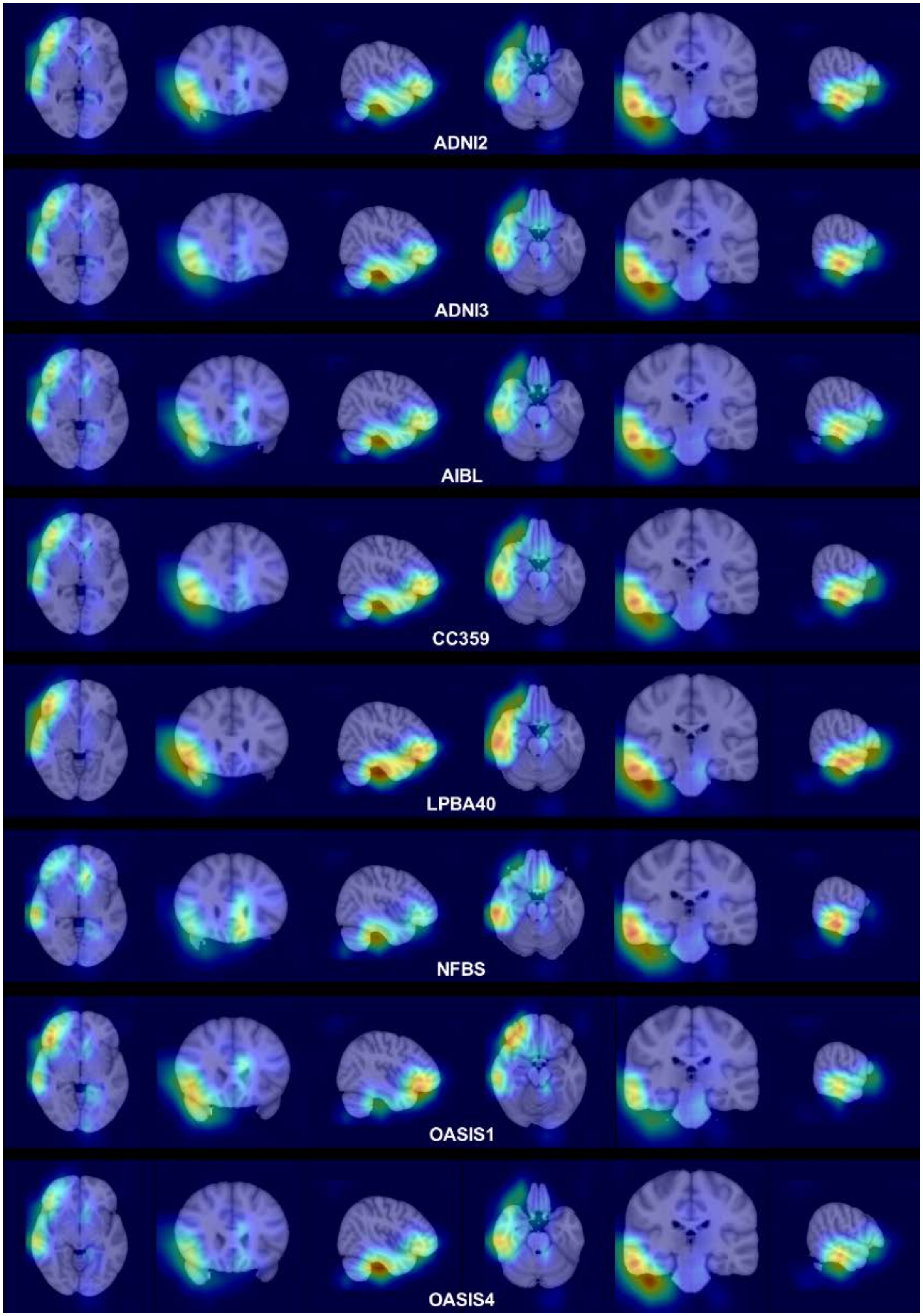

Supplementary Figure B. Average Grad-CAM per dataset in seed 1.

### Supplementary Figure C

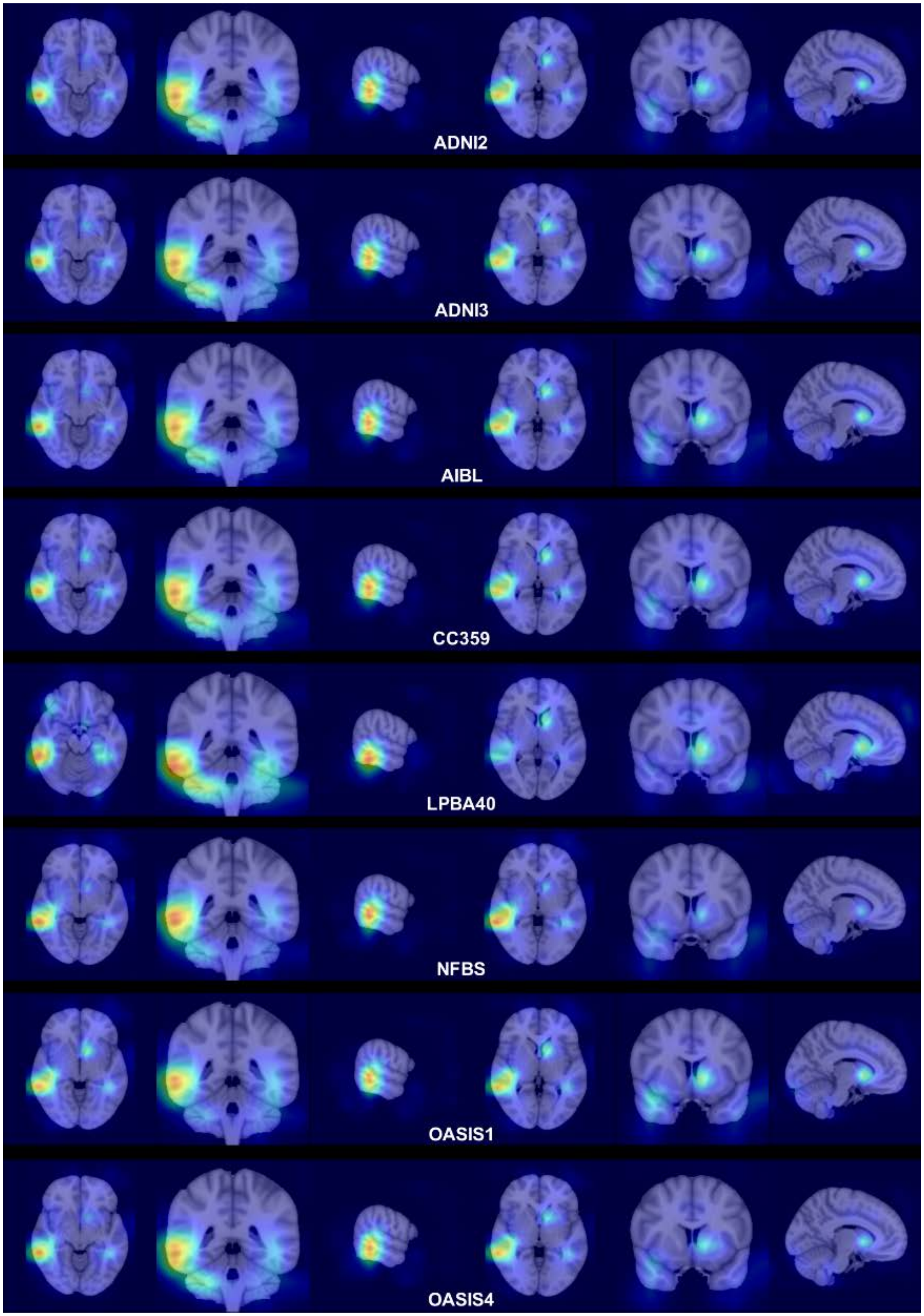

Supplementary Figure C. Average Grad-CAM per dataset in seed 2.

### Supplementary Figure D

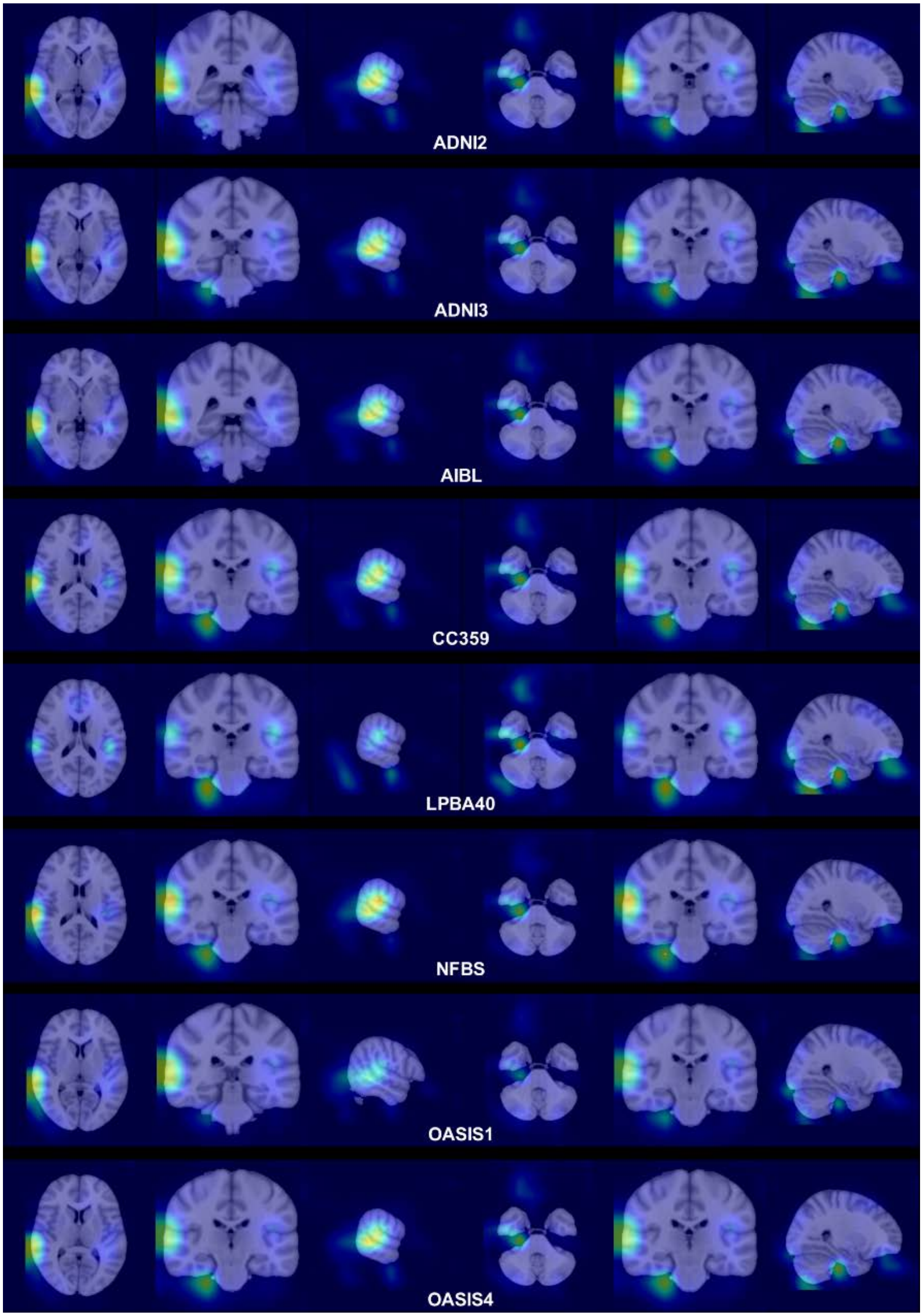

Supplementary Figure D. Average Grad-CAM per dataset in seed 3.

### Supplementary Figure F

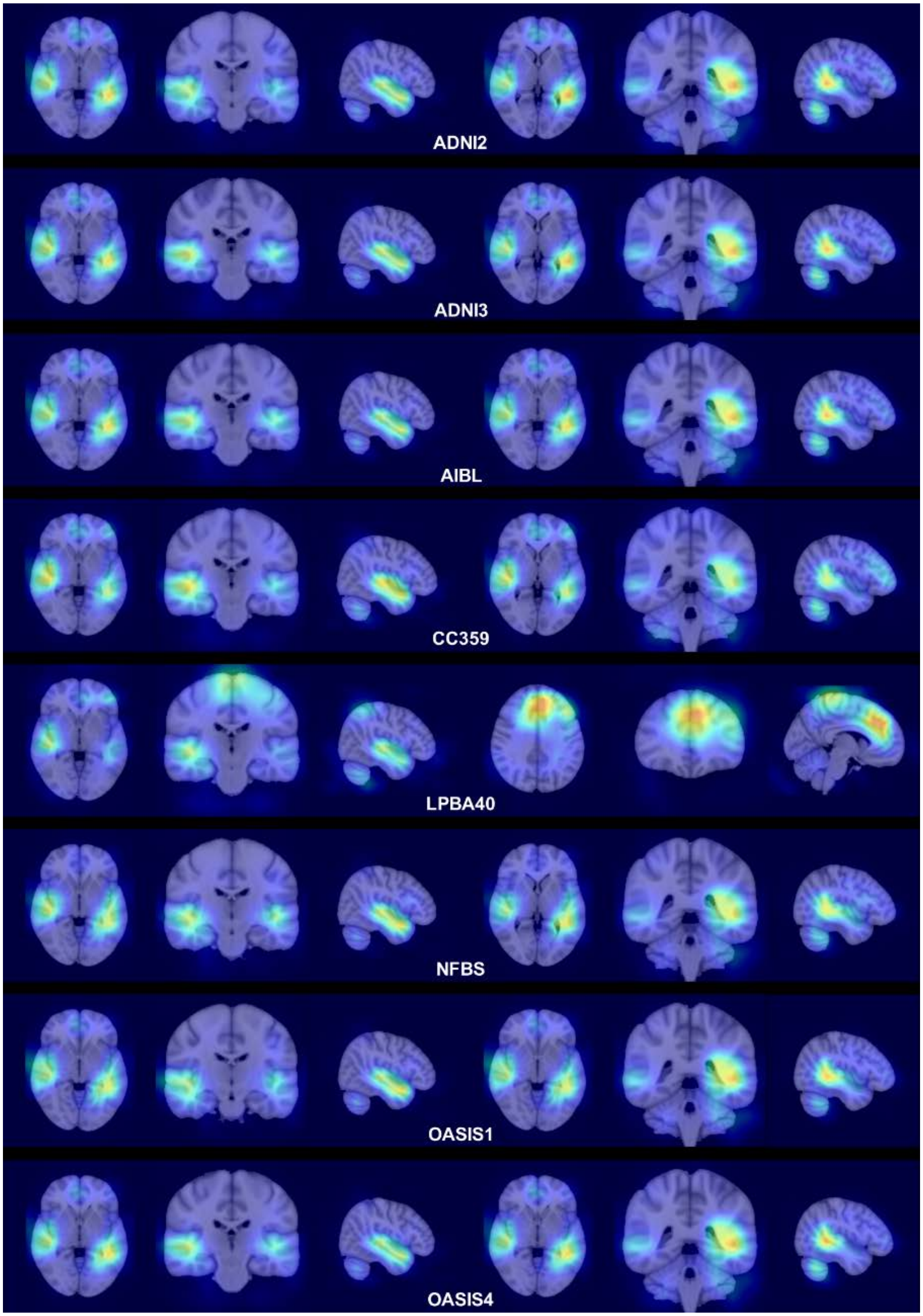

Supplementary Figure F. Average Grad-CAM per dataset in seed 5.
