## Supplementary Figure E for "A Neural Network Approach to Identify Left-Right Orientation of Anatomical Brain MRI"

Supplementary Figure E. Average Grad-CAM per dataset in seed 4.

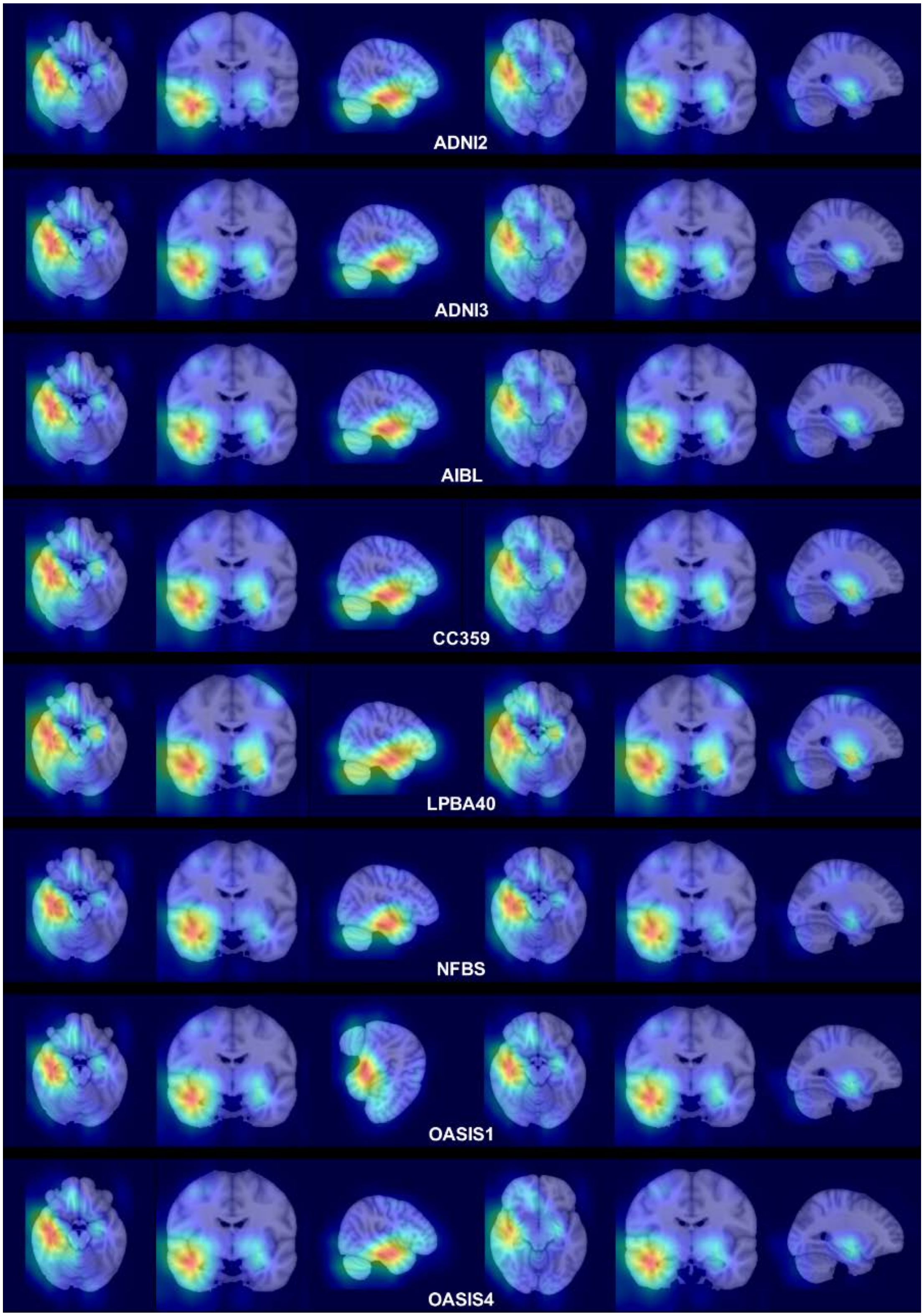


Supplementary Figure E. Average Grad-CAM per dataset in seed 4.
